# Light-sheet fluorescence microscopy of optically cleared human head-and-neck organs: three-dimensional tissue assessment and histological correlation

**DOI:** 10.64898/2026.10.30.763209

**Authors:** Lisa D.J. Schiffelers, Mari F.C.M. van den Hout, Ioana Gabor, Albert Bitorina, Elias Wieland, Taisiya Ageeva, Andreas Herrler, Zohaib Salahuddin, Francesco Missale, Anna Schueth

**Affiliations:** Department of Genetics & Cell Biology, Maastricht University (UM), Maastricht, the Netherlands; Department of Pathology, Maastricht University Medical Centre (MUMC+), Maastricht, the Netherlands; Department of Anatomy & Embryology, UM, Maastricht, the Netherlands; Department of Precision Medicine, UM, Maastricht, the Netherlands; Department of Otorhinolaryngology, Head & Neck Surgery, MUMC+, Maastricht, the Netherlands; GROW Research Institute for Oncology and Reproduction, UM, Maastricht, the Netherlands; NUTRIM Research Institute for Nutrition and Translational Research in Metabolism, UM, Maastricht, the Netherlands

**Keywords:** light-sheet fluorescence microscopy, tissue clearing, head-and-neck cancer, palatine tonsil, tongue, parotid gland, cervical lymph node, volumetric imaging, haematoxylin and eosin

## Abstract

Conventional histology provides detailed cellular information but only a partial view of the three-dimensional (3D) organisation of head-and-neck tissues. We investigated whether optical tissue clearing combined with light-sheet fluorescence microscopy (LSFM) could visualise the microanatomy of large head-and-neck specimens and relate volumetric features to conventional histology. Non-neoplastic human palatine tonsil, cervical lymph node, tongue and parotid gland specimens were cleared with a solvent-based protocol, labelled with small-molecule dyes and imaged by LSFM, with haematoxylin and eosin (H&E) histology as a morphological reference. LSFM visualised the characteristic architecture of each tissue, including lymphoid follicles and crypt epithelium in tonsil, cortex, paracortex and follicles in lymph node, epithelium and intrinsic muscle fibres in tongue, and lobules, ducts and adipocytes in parotid. Following structures through successive optical sections revealed their continuity and branching over long distances, information not available from single sections. Tongue epithelial measurements were comparable with H&E, and segmentation enabled volumetric quantification of structures such as the tongue epithelium and parotid adipocytes. Optical clearing and LSFM thus provide a feasible single workflow for volumetric investigation of human head- and-neck microanatomy. These data establish a 3D morphological reference with potential applications in cancer research and staging, computational image analysis and anatomical education, to be further developed through larger donor cohorts and specific labelling.

## Introduction

The head-and-neck region is an anatomically complex area, consisting of various organs, such as the tongue, the larynx, the tonsils, the thyroid, and salivary glands like the parotid and submandibular gland. In these organs, epithelial, glandular, muscular, cartilaginous and lymphoid tissues are closely associated with vascular and neural networks(1). This complex three-dimensional (3D) tissue structure supports important functions including phonation, swallowing, respiration, secretion and immune surveillance.

To date, tissue slicing and conventional histopathology, paired with classic two-dimensional (2D) microscopy, remain the reference methods for evaluating tissue morphology in the academic and clinical settings. Haematoxylin and eosin (H&E)-stained tissue sections provide detailed cellular information but represent merely a small fraction of the 3D organisation of a larger tissue specimen. Even though reconstruction of large tissue volumes from serial sections can extend this view into 3D, this is very labour-intensive and susceptible to tissue distortion, section loss, and registration errors (2). As a result, many structural relationships extending across hundreds of micrometres to several millimetres cannot be readily appreciated using conventional 2D microscopy. Limited sampling can affect the assessment of tissue composition and spatial heterogeneity in both non-neoplastic and neoplastic tissues (3).

Volumetric imaging offers a complementary approach to examine structural continuity, thereby placing local histological features within their broader spatial context. Clinical imaging modalities such as MRI and CT enable 3D reconstruction of head-and-neck organs in vivo and support patient-specific biomechanical modelling (4–6), but at sub-millimetre resolution these approaches only capture organ geometry, without cellular or molecular information. Optical tissue clearing combined with light-sheet fluorescence microscopy (LSFM) provides a means of obtaining this detailed information from human tissue specimens. Clearing renders tissue samples transparent and reduces light scattering through refractive index matching by decreasing optical heterogeneity within the tissue. In this manner, large-scale millimetre to centimetre sized tissue samples can be assessed rapidly at high spatial resolution while preserving their native 3D architecture (2). Dedicated preparation and labelling workflows have enabled imaging of conventionally fixed human tissues and, following deparaffinisation, archived formalin-fixed, paraffin-embedded (FFPE) specimens (2,7). LSFM studies using tissue samples from various organs, such as prostate, brain, oesophageal, breast and lymph node have demonstrated recognisable microanatomical structures in comparison with conventional H&E sections, together with additional information on their spatial organisation (8–13). The combination of clearing, labelling, and 3D LSFM imaging has transformed volumetric imaging in different fields, such as developmental biology and neuroscience, and is increasingly being applied to human tissues for anatomical and pathological investigations.

Studies applying tissue clearing and 3D LSFM to head-and-neck tissues are emerging. One recent study reports the clearing of head-and-neck tumour biopsies from the tongue, cheek, and floor of the mouth, showing the promise of this approach for the application in a clinical diagnostic setting (14). Clearing protocols have also been evaluated for formalin-fixed human tonsil up to a thickness of 200 µm, with a focus on the visualisation of lymphoid tissue (15). Recently, CUBIC-cleared human tongue biopsies (approximately 2 x 3 x 3 mm) enabled multiplex whole-mount labelling of blood vessels, muscle fibres and nuclei (16). Other clearing and LSFM studies have already enabled a cellular atlas of the embryonic and foetal development of the human head (17), as well as imaging of adult donor-derived thyroid tissue (18). Together, these studies show the potential of 3D imaging for head- and-neck tissues, but each study addresses solely a single organ, a specific disease, a developmental stage, or a technical question. Moreover, these experiments were mostly performed on small biopsies of a few millimetres or on slices of only a few hundred micrometres thick, rather than on whole organs or large tissue specimens. In addition, imaging was not consistently performed with LSFM; several studies relied partly or entirely on confocal microscopy, which limits especially the acquisition speed. A systematic 3D reference of the normal adult human head-and-neck organs, imaged with a uniform workflow and directly correlated with conventional H&E histology, is still lacking.

Here, we present high-resolution volumetric LSFM mapping of optically cleared non-neoplastic human head-and-neck specimens obtained from donated cadavers. Our LSFM study comprises the palatine tonsil, cervical lymph node, tongue, and parotid gland. Large tissue specimens are cleared, labelled and imaged with a single pipeline, annotated with the compartments familiar from 2D histology, and visualised in both 2D and 3D. Our primary aim is to assess whether the principal microanatomical structures characteristic of each tissue can be recognised in LSFM volumes, using conventional H&E sections as a morphological reference. We further aim to explore selected aspects of their 3D organisation through volumetric visualisation, segmentation and reconstruction of structures of interest. In non-neoplastic head-and-neck organs, 3D imaging enables the characterisation of lymphoid structures, glandular architecture, ductal branching, vascular networks, epithelial boundaries, and the spatial relationships between these structures and surrounding connective tissue, providing insight into their organisation and continuity that may be difficult to appreciate in isolated, thin histological sections (17,19). Mapping these tissues across organs provides a morphological reference for interpreting volumetric images, and a resource for anatomical studies, education, quantitative morphometric analyses, and the development and validation of image analysis algorithms.

Because abnormal architecture is best recognised against a defined baseline, this dataset also establishes a reference against which pathological alterations, such as tumour invasion, inflammatory changes and lymphatic remodelling, can be evaluated in 3D. This work thereby bridges conventional histopathology and volumetric imaging, and provides a framework for future applications in anatomy, pathology and computational biomedical imaging.

## Results

### Sampling, clearing and labelling of human head-and-neck tissues

To study head-and-neck tissue samples, a diverse set of tissues was obtained from human donor cadavers and sliced in multiple parts for 2D H&E and 3D LSFM analyses. The cervical lymph node and the palatine tonsil were longitudinally cut in half, whereas the parotid gland and tongue were transversally cut in multiple pieces of up to 1.5 cm in thickness (Supplementary Figure 1). Tissues meant for light-sheet microscopic analysis were subsequently cleared using our iDISCO-like MASH protocol (11–13) and stained with small-molecule dyes (Table 1), which rendered all specimens transparent (Figure 1). All organs were imaged in three channels: neutral red (NR) fluorescence, and autofluorescence at 488 nm (AF488) and 640 nm (AF640) excitation. NR is a basic cationic dye that stains basophilic structures such as nucleic acids, whereas the autofluorescence channels provide complementary contrast of tissue components such as muscle and connective tissue. Each specimen was first imaged in its entirety at low magnification to provide a whole-sample overview, after which selected regions of interest were imaged at higher magnification and compared with H&E-stained sections from the matching tissue. In the following sections, we report on the 3D microanatomical architecture of each of the optically cleared and fluorescently labelled human head-and-neck organs.

**Figure 1.**
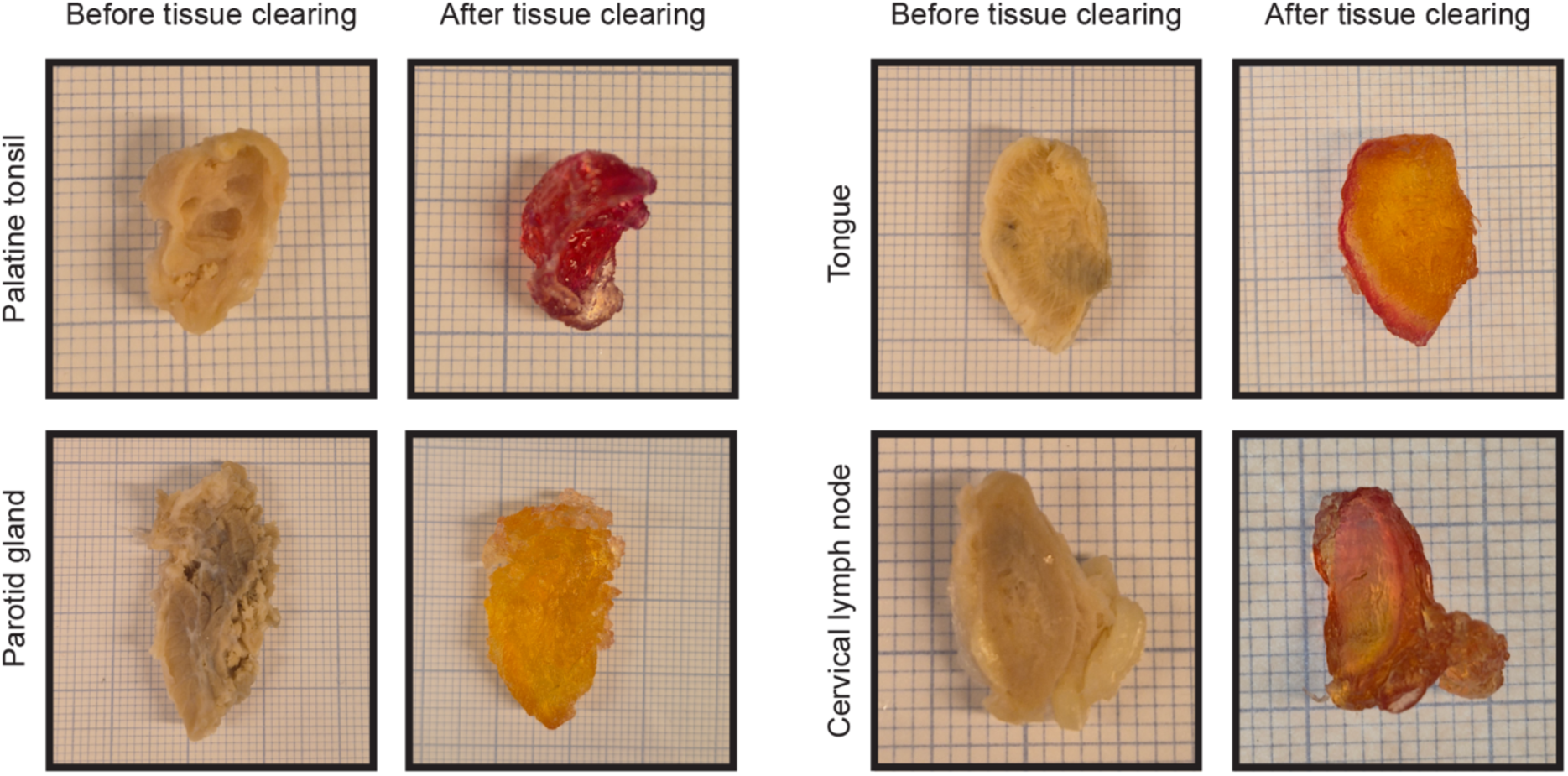
Human head-and-neck tissues before and after MASH clearing and labelling. Images of a palatine tonsil, a cervical lymph node, a tongue and a parotid gland are captured on millimetre paper before and after tissue clearing. Small square: 1×1 mm.

**Table 1.** Overview of fixed human cadaver tissues used for light-sheet microscopy imaging.

| **Organ** | **Dimensions pre-clearing (mm)** | **Staining** | **Channels recorded** |
|---|---|---|---|
| Palatine tonsil | 14 x 11 x 10.2 | DAPI+NR | 488 (AF), 561 (NR), 640 (AF) |
| Cervical lymph node | 14 x 11 x 5.8 | DAPI+NR | 488 (AF), 561 (NR), 640 (AF) |
| Tongue | 22 x 13 x 7.8 | DAPI+NR | 488 (AF), 561 (NR), 640 (AF) |
| Parotid gland | 37 x 20 x 12.2 | DAPI+NR | 488 (AF), 561 (NR), 640 (AF) |

AF, autofluorescence; DAPI, 4’,6-diamidino-2-phenylindole; NR, neutral red. DAPI (800 nm) was only recorded by two-photon microscopy (TPM) (parotid gland, Supplementary Figure 6).

### 3D imaging of the palatine tonsil reveals lymphoid follicles, crypt epithelium and tubular structures

For the 3D mapping of tonsil tissue, one half of a palatine tonsil (14 x 11 x 10.2 mm) was cleared, labelled, and imaged by LSFM. (Figure 2A,B). Imaging through the entire tissue enabled visualisation of the architecture of the imaged volume of 1.57 cm^3^ in cross-sectional 2D sections and a full 3D overview (Figure 2A,B; Supplementary Figure 2A). Selected regions of interest were subsequently imaged at higher magnification to examine their structural and cellular organisation in greater detail. A side-by-side comparison with H&E-stained sections revealed similar tissue characteristics. In the epithelial layer, structures such as the epithelial-connective tissue interface and the deep invaginations into the lymphoid tissue, also known as tonsillar crypts, can be observed (Figure 2C-E). Furthermore, several lymphoid follicles could be detected, and the follicular cells could be distinguished from the interfollicular area and visualised throughout the volume of the palatine tonsil (Figure 2C,E; Supplementary Figure 2B). Imaging the tonsil sample at the highest magnification of 30× enabled clear visualisation of fine tissue structures while maintaining high image quality (Figure 2F). Individual nuclei remained distinguishable, and individual cells lining the tubular structures could be discerned. Diameters of these tubular structures, that most likely represent blood vessels, could also be measured (e.g. 97 and 139 µm), revealing variation in vessel calibre and enabling visualisation of both larger tubules and smaller branches. Furthermore, visualisation of the tubular network in orthogonal views enabled tracing through the tissue volume, as exemplified by a tubular structure followed over approximately 130 µm (Supplementary Figure 2C,D). These observations suggest that three-dimensional reconstruction of the lymphovascular network is feasible with targeted labelling.

**Figure 2.**
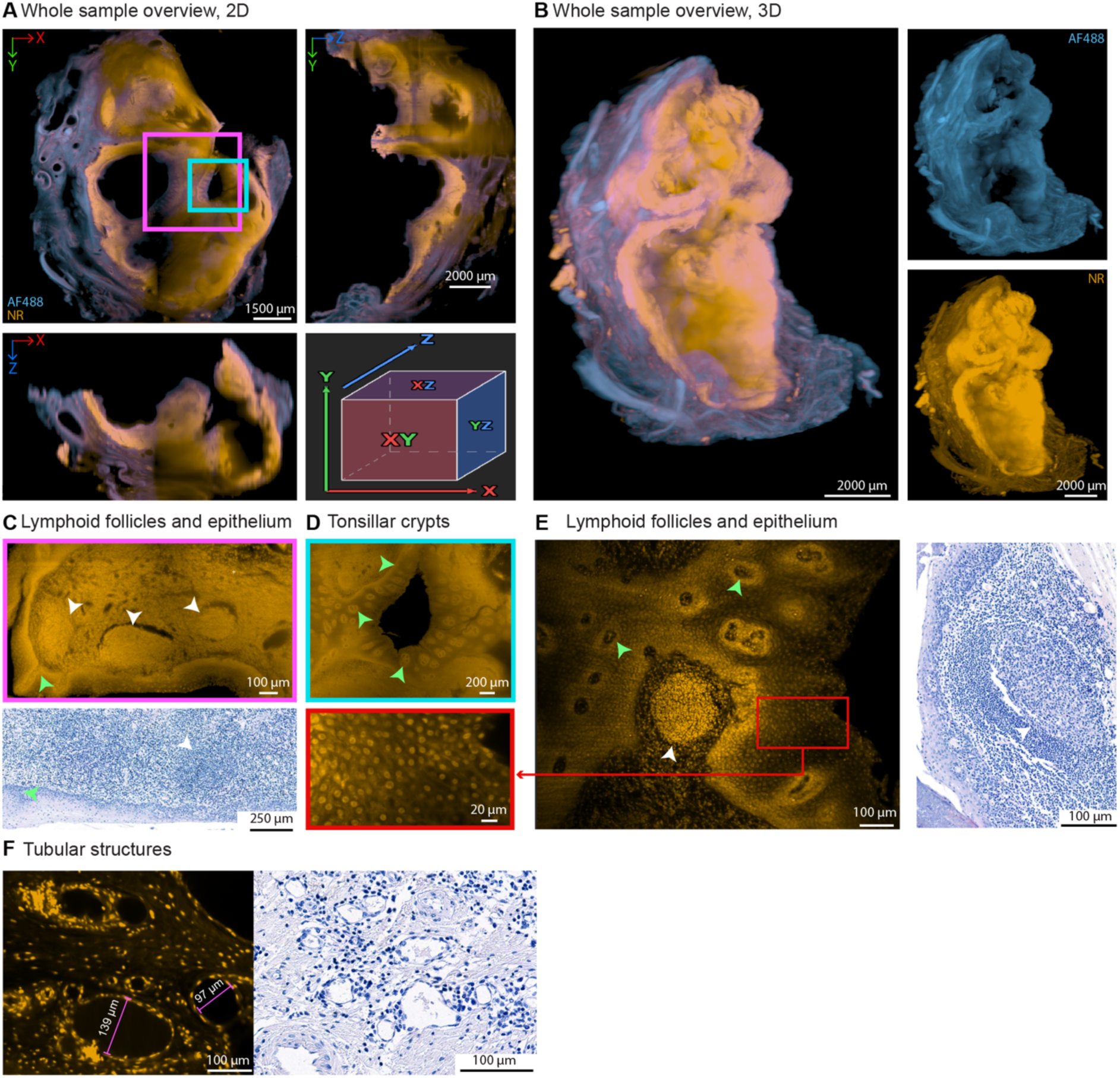
3D architecture of the palatine tonsil with its lymphoid follicles, crypt epithelium and tubular structures. One half of the tonsil was cleared, labelled with neutral red (NR) and imaged by LSFM; the other half was processed for H&E. **A)** 2D whole sample overview including orthogonal views. The boxed regions are enlarged in C and D. **B)** 3D representation of the entire half of the tonsil tissue. **C-F)** Region of interest imaged at a higher magnification and compared to H&E-stained sections to visualise lymphoid follicles (white arrowheads) (C, E), tonsillar crypts in the epithelium (green arrowheads) (C, D, E), epithelial cells (E, including the boxed red region), and tubular structures (F) in greater detail. Pseudocolours: autofluorescence (AF)488 (blue), NR (yellow). Scale bars and their measures are depicted in the lower right corner of each panel.

### 3D imaging of a cervical lymph node maps its compartments and lymphoid follicles

One half of a bisected cervical lymph node, measuring 14 x 11 x 5.8 mm, was visualised with LSFM. In the 2D and 3D renderings of this imaged volume of 0.89 cm^3^, the outer cortex, the inner paracortex and the central medulla could be distinguished (Figure 3A,B; Supplementary Figure 3A). Autofluorescence (488 nm excitation) clearly delineated the hilum, located at the bottom of the lymph node, together with the hilar lymphovascular structures branching through its central region (Figure 3B). On the right side, lobulated perinodal adipose tissue was attached to the capsule. The virtual 2D sections from the LSFM volume showed the same macroscopic components as the corresponding large-scale overview H&E-stained sections (Supplementary Figure 3B). When focussing on the border of the lymph node cortex, primary follicles (B-cell zones), the paracortex (T-cell zone) and the capsule, the dense connective tissue layer forming the outer boundary, can be detected by both H&E and LSFM (Figure 3C). Interestingly, also secondary lymphoid follicles could be found with a germinal centre and mantle zone, which could give additional information regarding the immune status of the tissue (Figure 3D). The measured diameters of the primary follicles ranged from 88 to 347 µm (Supplementary Figure 3C). In addition, tubular structures that possibly represent lymph or blood vessels could be detected and traced through the LSFM volume, as exemplified by a tubular structure followed over approximately 320 µm (Figure 3E; Supplementary Figure 3D) and approximately 1500 µm (Supplementary Video 1), which is not possible on 2D H&E sections.

**Figure 3.**
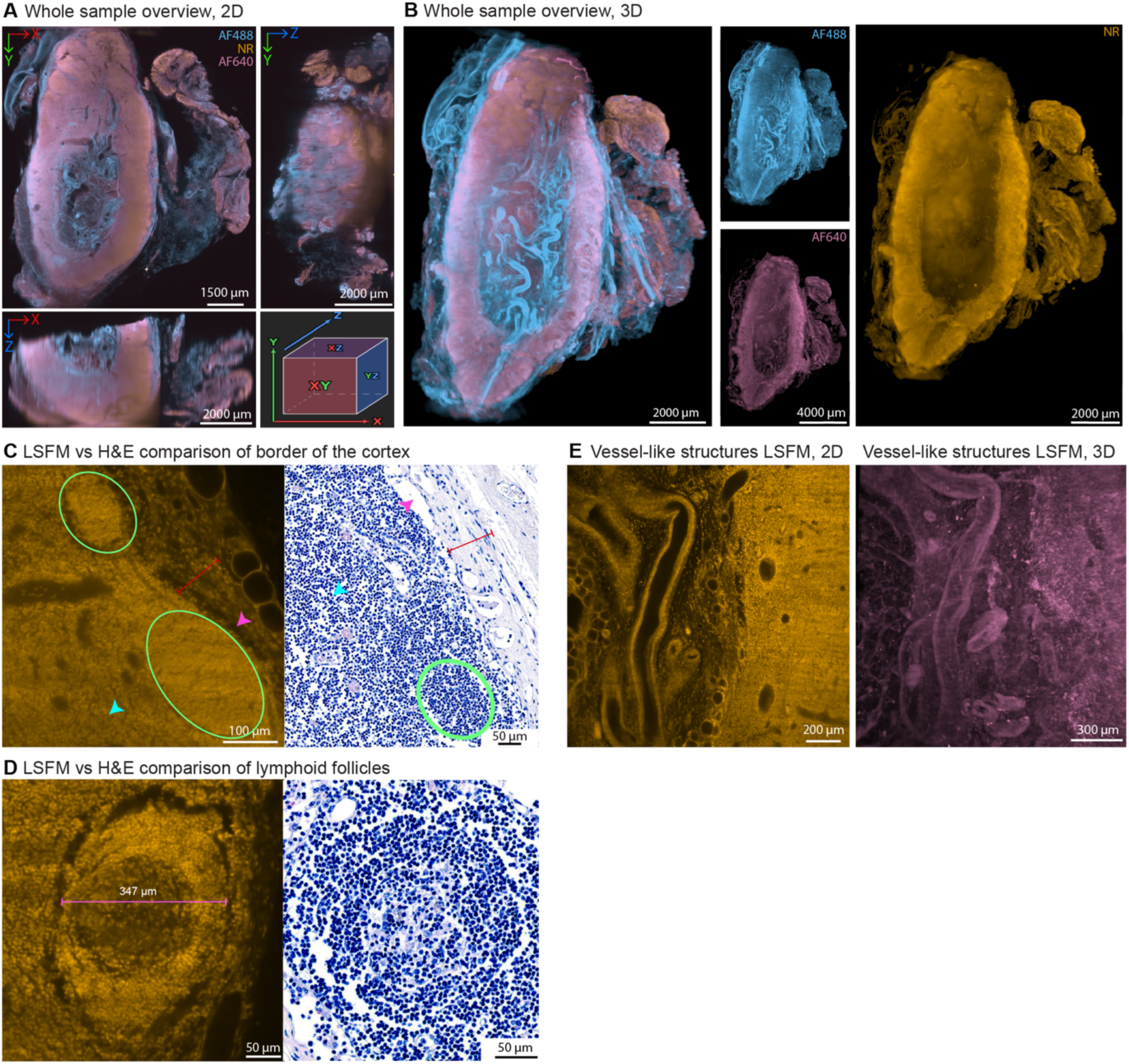
3D architecture of a cervical lymph node, including lymphoid follicles and tubular structures. One half of the lymph node was cleared, labelled with neutral red (NR) and imaged by LSFM; the other half was processed for H&E. **A)** 2D whole sample overview including orthogonal views from the indicated directions. **B)** 3D representation of the entire cleared lymph node tissue, including the separate channels. **C)** Region of interest imaged at a higher magnification with matching H&E images, to visualise the capsule (red line), subcapsular sinus (pink arrowheads), primary follicles (encircled in green) and the paracortex (blue arrows) in greater detail. **D)** Secondary lymphoid follicle in both LSFM and H&E modalities. **E)** Tubular structures in 2D and 3D LSFM representations. Pseudocolours: autofluorescence (AF)488 (blue), NR (yellow), AF640 (magenta). Scale bars and their measures are depicted in the lower right corner of each panel.

### 3D imaging of the tip of the tongue shows an intricate network of skeletal muscle fibres and tubular structures

Light-sheet microscopy of the tip of the tongue was performed on a tissue piece of 22 × 13 × 7.8 mm, as can be appreciated from the 2D cross-sectional views and the 3D overview of the full 2.2 cm³ volume (Figure 4A,B; Supplementary Figure 4A; Supplementary Video 2). Interestingly, the autofluorescence signals mainly captured the muscle architecture, whereas neutral red highlights the epithelial structure of the tongue (Figure 4B; Supplementary Figure 4A). A comparison of a region encompassing the epithelium, the connective tissue layer and the border of the intricate muscle network with H&E revealed a similar tissue architecture and comparable measures, with an epithelial thickness of 496 µm and 499 µm and a connective tissue thickness of 1110 µm and 1021 µm for LSFM and H&E, respectively (Figure 4C). It must be noted that the thickness of the epithelium and the connective tissue layer is highly diverse due to the presence of the different types of papillae and the rete ridges of the epithelium (Figure 4C; Supplementary Figure 4B). Hence, a volumetric quantification of the epithelium would be more meaningful as it is less dependent on the measured cross-section. Surface segmentation of the three-dimensional LSFM data set (with dimensions of 1849 x 1849 x 5628 µm) resulted in the extraction of the epithelial layer and measured a volume of 3.916 mm^3^ (Figure 4D). The 3D representation of this part of the tongue also showed the intricate network of lumen-containing tubular structures that protrude towards the epithelium, forming the basis for tubular segmentation and tracking analyses (Figure 4D). Even though blood vessels can also be clearly visualised in H&E, their continuity and branching pattern are difficult to assess on these thin sections (Figure 4E). LSFM could show that lumen-containing tubular structures with a cellular lining are often already connected as exemplified by their tracing to a branching point through consecutive optical sections over approximately 450 µm (Supplementary Figure 4C), which is information that could not be deduced from the single 4 µm H&E slice. With the right 3D analysis tools and markers, the whole vasculature, ductal system, and nerve pathways could be extracted and visualised as a complete network to give insights into the organisation of the whole tissue. The tongue also contains interwoven intrinsic muscle fibres oriented in longitudinal, transverse, and vertical directions. Upon zooming into these muscle layers, the transversal and longitudinal muscle fibres of the tongue can be clearly visualised up to a nuclear resolution (Figure 4F). The H&E only offers a planar view, but LSFM imaging throughout the complete thickness of 7.8 mm allowed individual muscle fibres to be followed through consecutive optical sections, as exemplified by fibres tracked over approximately 1500 µm (Supplementary Figure 4D). Subsequent surface segmentation enabled the recreation of the muscle fibres in 3D (Figure 4G).

**Figure 4.**
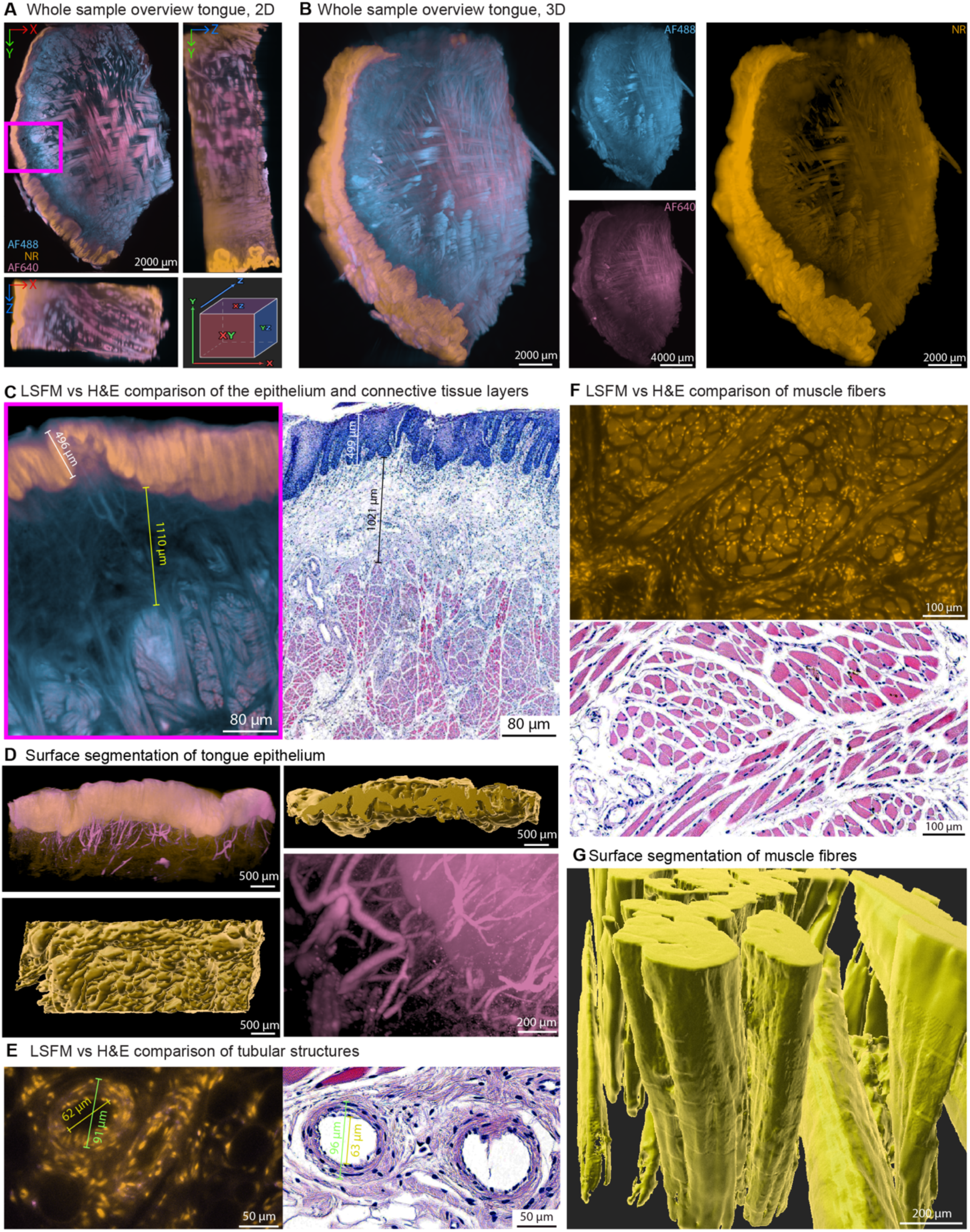
3D architecture of the tongue tip with its epithelium, tubular structures and intrinsic muscle fibres. A piece of the tongue tip was cleared, labelled with neutral red (NR) and imaged by LSFM; a separate piece was processed for H&E. **A)** 2D whole sample overview including orthogonal views from the indicated directions. The boxed region is enlarged in C. **B)** 3D representation of the entire tongue tissue, including the separate channels. **C, E, F)** Region of interest imaged at a higher magnification with matching H&E images, to visualise the epithelium and connective tissue layer (C), tubular structures (E), and the muscle network (F) in greater detail. **D, G)** Surface segmentation to extract three-dimensional structures of the epithelium (D) and the muscular fibres (G). Pseudocolours: autofluorescence (AF)488 (blue), NR (yellow), AF640 (magenta). Scale bars and their measures are depicted in the lower right corner of each panel.

### 3D imaging of the parotid gland captures its lobular organisation, ductal network and adipocytes

The parotid gland is a purely serous, branched acinar gland with abundant adipocytes and striated ducts(20). For LSFM, a parotid tissue piece of 37 × 20 × 12.2 mm was cleared, labelled and imaged (Figure 5A,B). Its lobular organisation can already be appreciated macroscopically in the 2D cross-sectional views of the light-sheet fluorescence imaging results and the H&E-stained tissue slice (Figure 5A; Supplementary Figure 5A,B). 3D visualisation of the complete 9.03 cm^3^ tissue sample showed comparable structures in the imaged channels but also demonstrated that the image quality decreased towards the centre of the tissue, where both resolution and fluorescence signal were reduced (Figure 5B). In the large-scale images of both the LSFM and the H&E, the connective tissue capsule can be observed, and its inward branching septa divide the parotid gland into clearly delineated lobules of 1-3 mm (Figure 5A,C). The serous acini first secrete their protein-rich secretion product into their own lumen, after which they drain into the intralobular ducts (21). Interlobular ducts, that run within the septa, then guide the secretion product towards the interlobar ducts and finally the parotid duct or Stenson’s duct (21). Interestingly, the tubular structures in the LSFM data are particularly visible in the 640 nm autofluorescence channel (Figure 5D; Supplementary Figure 5C). The volumetric dataset allowed the tracking of these intralobular ducts through consecutive optical sections, as exemplified by ducts followed over approximately 240 µm, and revealed that they branch from the same origin (Supplementary Figure 5C). In fact, a 3D view of one lobule demonstrated an organised and intricate tubular structure network (Figure 5E; Supplementary Figure 5D; Supplementary Video 3).

**Figure 5.**
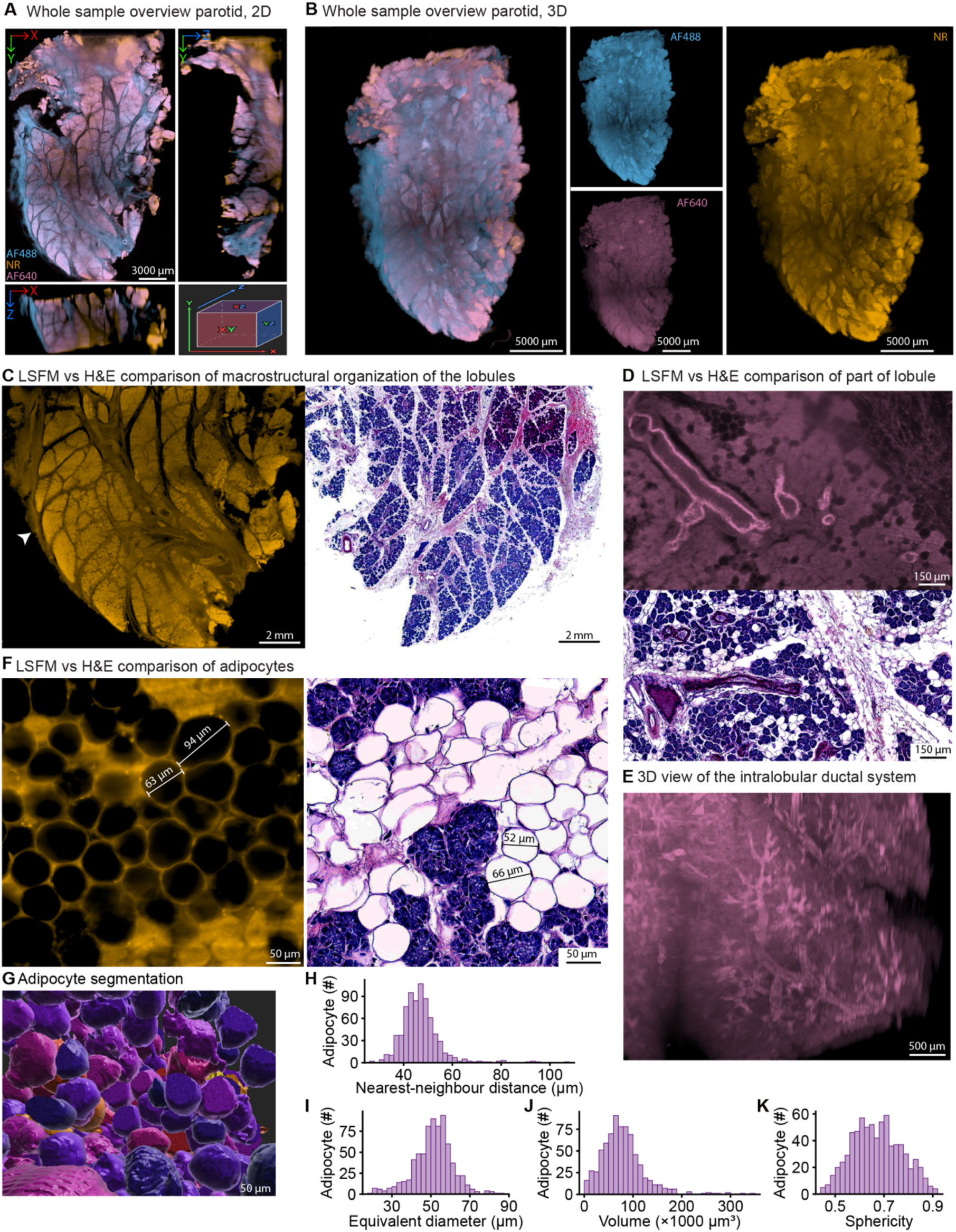
3D architecture of the parotid gland with its lobules, ductal network and adipocytes. A piece of the parotid gland was cleared, labelled with neutral red (NR) and imaged by LSFM; a separate piece was processed for H&E. **A)** 2D whole sample overview including orthogonal views from the indicated directions. **B)** 3D representation of the entire parotid gland tissue, including the separate channels. **C, D, F)** Region of interest imaged at a higher magnification with matching H&E images, to visualise the typical lobular organisation of the parotid with its capsule (white arrowhead) (C), intralobular tubular structures (D), and adipocytes (F) in greater detail. **E)** Three-dimensional representation of the intricate intralobular ductal system. **G)** Adipocytes upon surface segmentation. Individual adipocytes are shown in random colours for visual distinction only. **H-K)** Segmentation-based calculation of the nearest-neighbour distance (H), diameter (I), volume (J) and sphericity (K) of the adipocytes. For sphericity, 1 is a perfect sphere. Pseudocolours: autofluorescence (AF)488 (blue), NR (yellow), AF640 (magenta). Scale bars and their measures are depicted in the lower right corner of each panel.

Adipocytes are part of the stroma, and their numbers are known to increase with age (22,23). At higher magnifications, adipocytes could be visualised in greater detail (Figure 5F). Adipocyte segmentation was performed on an LSFM stack, covering a total volume of 0.38 mm³ (444 × 444 × 1928 µm) (Figure 5G; Supplementary Figure 5E). In this volume, 1191 adipocytes were identified, corresponding to a density of approximately 3.1 x 10^3^ adipocytes per mm^3^. Together, they occupied 22% of the imaged volume (Supplementary Figure 5F). Of these, 733 adipocytes were fully contained within the volume without touching the image border and were used for the size and shape measurements. The mean centroid-to-centroid distance to the nearest neighbouring adipocyte was 46.8 ± 8.1 µm (Figure 5H). This is close to the mean adipocyte diameter (Figure 5I), indicating that most adipocytes were in direct contact with a neighbour and thus occurred in local clusters. The adipocytes had an average diameter of 51.3 ± 10.1 µm, an average volume of 78539 ± 43838 µm^3^ and an average sphericity of 0.672 ± 0.100 (Figure 5I-K).

For an additional higher resolution, multi-modal comparison, we also performed two-photon microscopy (TPM) experiments on optically cleared parotid tissue. Similar structures such as adipocytes, acini, connective tissue and tubular structures, could be detected in the TPM imaging data at higher resolution (Supplementary Figure 6A). Following five consecutive optical sections at 10 µm intervals over a depth of 40 µm showed how the morphology of individual structures, such as ducts and acini, changed with imaging depth (Supplementary Figure 6B). TPM data were correlated with H&E-stained sections to identify structures and architectural features relevant to head-and-neck clinicians, with interpretation confirmed by an experienced pathologist. In addition, second harmonic generation visualised collagen-rich connective tissue without labelling, illustrating how TPM can complement LSFM with cellular detail in selected regions, whereas LSFM provides the overview of the complete specimen

## Discussion

This study demonstrates that solvent-based clearing, small-molecule labelling and light-sheet fluorescence microscopy (LSFM) can be applied to large human head-and-neck specimens of up to approximately 12 mm in thickness, reproducing the principal microanatomical features known from conventional histology. Across tonsil, cervical lymph node, tongue and parotid gland, a single workflow connected recognisable histological features with their three-dimensional (3D) organisation. The principal advantage of cleared-tissue LSFM on head-and-neck tissue samples is the ability to follow structures continuously within large volumes, preserving spatial relationships that are only partially represented in individual sections, while selected segmentations showed that these volumes are directly amenable to quantitative analysis. Haematoxylin and eosin (H&E) histology remained an essential reference, and the close correspondence between both modalities supports the use of LSFM volumes as a morphological reference for head-and-neck tissues.

Previous studies established clearing and volumetric imaging of large human brain and prostate samples (12) and even of intact adult human organs (18). In the head-and-neck region, studies have so far focused on biopsies of a few millimetres or slices of a few hundred micrometres (14–16). Our findings extend this work to structurally diverse head-and-neck tissues with volumes approximately two orders of magnitude larger, allowing structures to be followed over long distances, compartments to be studied in relation to each other, and regional heterogeneity to be assessed. These volumes also complement MRI- and CT-based head-and-neck atlases (5), which capture organ geometry but not microscopic tissue architecture. The parotid gland was the largest cleared and imaged specimen (37 × 20 × 12.2 mm). Its lobular organisation remained visible throughout, while signal and image quality decreased towards the centre, defining the current practical range of the workflow. Clearing, dye penetration and optical performance can be optimised separately for specimens of this size.

Tonsillar and lymph-node reconstructions provide a framework for examining follicular number, volume, shape, nuclear density and distribution within surrounding compartments. Following follicles through depth reduces misclassification of primary and secondary follicles when a germinal centre lies outside an individual section, and whole-volume follicle counts could describe the state of the adaptive immune response more completely than sampling. Because cervical lymph nodes are the main site of regional spread of head-and-neck squamous cell carcinoma, these volumes may also serve as a baseline for 3D detection of occult metastases, whose size is systematically underestimated in 2D sections (24). The clearly delineated capsule moreover suggests that 3D assessment of extranodal extension, an adverse prognostic factor in the tumour-node-metastasis (TNM) staging, is feasible (25,26). B-cell and T-cell markers would improve delineation of follicles and paracortex, as illustrated by multiplex imaging of human tonsils and reactive lymph nodes (27), and endothelial or elastic-fibre labelling would separate the efferent lymphatic from the blood vessels in the hilum.

Most head-and-neck cancers arise in the epithelium, making it a key target for 3D reference data. Because epithelial thickness varies with crypts, papillae and rete ridges, volumetric measures such as epithelial volume and local thickness are more reliable than single sections. In the tonsil, a 3D map of the crypt network is particularly relevant, as human papillomavirus (HPV)-associated oropharyngeal carcinoma often arises from the crypt epithelium, where early lesions can remain hidden (28,29). In the tongue, a 3D reference of the epithelial to connective tissue interface could support detection of dysplasia (30), assessment of depth of invasion for TNM staging (26) and, ultimately, inspection of resection margins, which guides the treatment intensification in head-and-neck cancer (31). Beneath the epithelium, the interwoven intrinsic muscle fibres of the tongue could be followed through the volume and reconstructed in 3D. At cellular resolution, such data could serve as a reference for in vivo diffusion tensor tractography of tongue myoarchitecture (32), which is relevant for the investigation of swallowing disorders.

In the parotid, 3D imaging revealed lobular organisation and branching tubular structures, although distinguishing duct types, such as intercalated and striated ducts, remained easier in H&E. Specific epithelial and endothelial labelling would allow ducts and vessels to be identified and reconstructed as separate networks. Moreover, the adipocyte composition of the parotid gland is a relevant characteristic, as glandular fat content increases with age and body mass index (22,23) and in Sjögren syndrome (33). Whereas MRI fat-fraction mapping reports fat content per voxel, our segmentation of 1191 adipocytes resolved their number, size, shape and clustering, and could serve as a microscopic reference for such imaging biomarkers. Measuring directly in 3D also avoids a limitation of 2D morphometry, in which profile diameters depend on the height of the section through each cell (34). Of note, adipocyte profiles in our H&E sections were frequently distorted and unsuitable for segmentation, but it remains to be determined if this reflects tissue preservation or sectioning artefacts.

Altogether, these findings point to promising applications of volumetric imaging in head-and-neck anatomy, pathology and cancer research. However, some limitations should be considered. Each organ was represented by a single specimen from donor tissue that had been formalin-fixed for several years, so the measurements should be regarded as illustrative rather than as normal values. In addition, preparation for both LSFM and H&E may cause some shrinkage, image quality decreased with depth in the largest specimen, and the small-molecule dyes used here visualise architecture rather than specific cell types. Larger studies across donors, combined with specific labelling and independently validated segmentation, will be needed to establish reproducibility and normal ranges.

Future applications include quantitative computational image analysis of expert-annotated volumes. In prostate cancer, deep-learning analysis of 3D pathology has already yielded informative architectural features (35), outperformed 2D approaches in predicting recurrence and enabled triage of high-risk cross-sections for pathologist review (36,37). Normal 3D architecture also provides the baseline against which tumour invasion and immune infiltrates can be recognised, as illustrated by 3D mapping of T cells in head-and-neck cancer (38). Navigable volumes could further help learners relate familiar sectional appearances to their surrounding anatomy, and faster protocols (14) will be key to clinical translation.

In conclusion, optical clearing combined with LSFM enables volumetric visualisation of large human head-and-neck specimens across lymphoid, epithelial, muscular and glandular tissues, providing a 3D morphological reference and a foundation for quantitative and diagnostic 3D pathology in this anatomically complex region.

## Material and methods

### Human tissue samples

Human cadaver tissues from the head-and-neck region were obtained from donors enrolled in the body donation program of the Department of Anatomy & Embryology at Maastricht University (Maastricht, the Netherlands). The tissue donors gave their informed and written consent to the donation of their body for teaching and research purposes as regulated by the Dutch law for the use of human remains for scientific research and education (“Wet op de Lijkbezorging”). A handwritten and signed codicil, drawn up by each donor when still alive and well, is kept at the Department of Anatomy & Embryology. The bodies were preserved according to the department’s standard formalin-based fixation protocol and stored for 3 to 5 years. An overview of the samples derived from four different organs, dissected by an experienced head-and-neck surgeon, can be found in Table 1. Where possible, a matching piece of each organ was cut for the generation of a formalin-fixed, paraffin-embedded (FFPE) tissue block. Afterwards, the tissue samples were stored in 4% paraformaldehyde until further processing.

### MASH tissue clearing and labelling

All human cadaver head-and-neck tissues were subjected to our previously published iDISCO-like MASH protocol (11–13). Briefly, the fixed tissue samples were dehydrated using an increasing concentration of methanol, after which the samples were bleached overnight in 5% hydrogen peroxide to reduce pigmentation and autofluorescence. After rehydration, the tissues were permeabilised for 2 days in 0.2% Triton X-100 and stained for 7 days using two small-molecule fluorescent dyes: DAPI (4ʹ,6-diamidino-2-phenylindole; 1 µg/mL) as a nuclear counterstain for TPM and neutral red (NR) (0.001% [w/v]; 10 µg/mL) to visualise the basophilic (acidic) structures such as nucleic acids with LSFM. Following staining, the samples were dehydrated once more and delipidated using a 1:3 methanol in dichloromethane (DCM) solution to reduce light scattering. Ethyl cinnamate (ECi) was used for refractive index (RI) matching (RI ≈ 1.56) and served as the storage solution until light-sheet microscopy imaging. A step-by-step guide is described in the recently published bio-protocol by Schiffelers et al (11). Importantly, to optimise dye penetration into our relatively thick head-and-neck tissues (up to 12 mm), we prolonged permeabilisation and staining times to the above-mentioned durations.

### Generation of FFPE blocks

To enable a side-by-side comparison of our 3D images to the routine pathology pipeline, FFPE blocks were generated at the Department of Anatomy & Embryology. The matching tissue pieces of the organs listed in Table 1 were rinsed in Milli-Q water and subjected to a graded ethanol series (50%, 70%, 80%, 90%, 96%, and 100% (2x)) for dehydration, followed by two incubation steps in 100% xylene for clearing. Each incubation step was performed for 1 h per mm of tissue thickness. The tissues were subsequently transferred to molten paraffin with a melting temperature of 54-56°C and subjected to an overnight vacuum for the proper infiltration of paraffin into the tissue. FFPE block generation was performed using a Medite embedding device and a Tissue-Tek cooler. Afterwards, the blocks were stored at room temperature.

### Haematoxylin and eosin (H&E) staining

The generated FFPE blocks were cut into thin slices of 4 µm at the Department of Pathology, using a Jung Multicut 2045 rotary microtome (Leica Microsystems, Wetzlar, Germany). Single slices were then mounted on glass slides and allowed to dry. H&E staining was performed according to a routine histology protocol, also at the Department of Pathology. Briefly, sections were deparaffinised in xylene (1 min, 2x) and rehydrated through a graded ethanol series: 100% ethanol (1 min, 2x), 96% ethanol (1 min), 70% ethanol (1 min), and 50% ethanol (1 min). Following a brief rinse (10 s, 3x) in demineralized water, the sections were stained for 1 min in haematoxylin (solution modified according to Gill II, Sigma-Aldrich), followed by a 10 min rinse in tap water. Sections were subsequently stained with 0.5% (w/v) eosin (Sigma-Aldrich) (2 min) and rinsed again in tap water. The thin tissue slices were then dehydrated through an increasing ethanol series: 70% ethanol (1 min), 96% ethanol (1 min, 2x), and 100% ethanol (1 min, 2x). Finally, sections were cleared in xylene (1 min, 2x) and slides were mounted with xylene for long term storage using the Tissue-Tek SCA (Sakura) automated mounting system. Tissue-Tek® coverslipping film (Sakura) was used to seal the slides. Whole-slide scans were acquired using the Pannoramic 1000 scanner (3DHISTECH, Budapest, Hungary) with a 40x (NA 0.95) objective. Slide scans were visualised using QuPath (v0.6.0)(39). For the interpretation of the H&E data, an experienced clinical pathologist was consulted.

### Light-sheet fluorescence microscopic assessment

Cleared and stained human head-and-neck tissue samples were imaged by light-sheet fluorescence microscopy (LSFM) using an UltraMicroscope Blaze (Miltenyi Biotec, Bielefeld, Germany) controlled by ImSpector Software (version 8.0.3). Samples were placed in an adjustable sample holder and submerged in approximately 500 mL ethyl cinnamate (RI = 1.558) for RI matching. The RI of the imaging medium was set to MACS IS 1.558 in the liquid settings section of the microscope software. Depending on tissue dimensions and the region of interest, imaging was performed using the 1.1x (NA 0.1), the 4x (NA 0.35), or the 12x (NA 0.53) objective, which, combined with the zoom factors listed below, resulted in total magnifications of approximately 0.66x to 30x. Fluorescence was excited using fixed laser lines at 488, 561, and 640 nm, with corresponding emission filters of 525/50, 620/60, and 680/30 nm, respectively. NR fluorescence was acquired using 561 nm excitation. Tissue autofluorescence was recorded using 488 nm excitation (AF488) and 640 nm excitation (AF640). The stain and channels acquired per tissue are listed in Table 1. Emitted fluorescence was captured using a 4.2-megapixel sCMOS camera with a pixel size of 6.5 × 6.5 µm.

Image acquisition was performed in Multi Color 3D mode using the Light-Speed mode, with a light-sheet thickness of 3.9 µm. Z-stacks were recorded either across the entire tissue sample or within defined regions of interest, with a constant Z-step of 3 µm. This step size is slightly smaller than the light-sheet thickness of 3.9 µm, so consecutive optical sections overlapped, and no tissue was left unsampled between planes. Other acquisition parameters, including laser power, exposure time, zoom, and tiling options, were adjusted for individual samples according to tissue dimensions, staining intensity, region of interest, and signal-to-background ratio. Laser power software settings ranged from 5-60%, exposure times from 20-80 ms, and zoom factors from 0.6-2.5x. Where applicable, tiled image acquisition was performed in the Multi Color Mosaic Acquisition mode to cover the entire tissue or larger regions of interest. The resulting image stacks were exported as single-plane TIFF files and, additionally, the MACSiQView 3D Large Volume software (version 1.2.0) allowed for the generation of an Imaris file (.ims) and, wherever necessary, the automated stitching of the separate tiles.

### Two-photon microscopy

Cleared and stained human parotid gland tissue was imaged by two-photon microscopy (TPM) using an FV4000MPE multiphoton laser scanning microscope (Evident Scientific, Hamburg, Germany) controlled by CellSens FV software (version 4.1.1.137). The sample was placed in a Petri dish, and a few drops of ethyl cinnamate (RI = 1.558) were added for RI matching. A glass coverslip was placed on top of the sample to separate the cleared tissue from the water used as immersion medium for the 25x water-immersion objective (NA 1.05). Second harmonic generation (SHG) and DAPI fluorescence were excited at 800 nm (1% laser power) and detected using emission filters of 395/11 nm and 431/63 nm, respectively. The SHG channel visualised collagen-rich connective tissue, and the DAPI channel visualised nuclei. Images were acquired at 1024 × 1024 pixels with a pixel dwell time of 40 µs and a Z-step of 5 µm. TPM image stacks were analysed in Fiji (version 2.16.0)(40). Individual structures, such as ducts and acini, were tracked manually through consecutive optical sections to follow changes in their morphology with imaging depth (Supplementary Figure 6B).

### Image processing and analysis

Acquired LSFM data sets were analysed in Imaris (version 11.0.1, Oxford Instruments, Zurich, Switzerland). Imaris was used for three purposes: 1) the generation of 2D and 3D overview images, detailed images of regions of interest and videos, 2) manual measurements and tracking of structures through the image volume, and 3) surface segmentation for the volumetric quantification of tissue structures. The analyses are described below in the order in which they appear in the figures. LSFM stacks were imported as Imaris files (.ims). H&E whole-slide scans (.mrxs) were viewed and measured in QuPath. Segmentation statistics were exported from Imaris as .csv files for further calculations (see Statistics).

### Visualisation and manual measurements

Whole-sample overviews were generated as 2D optical sections with orthogonal (XZ and YZ) views (Figures 2A–5A; Supplementary Figures 2A–5A), and 3D overviews of the complete tissue samples as volume renderings in the Imaris 3D view (Figures 2B–5B). Regions of interest acquired at higher magnification were displayed in the same way and compared with H&E images of the corresponding region in the matching FFPE block. Supplementary Videos were created with the Imaris animation tool and exported as .mp4 files, and figure panels were exported as .tif snapshots. Brightness and contrast were adjusted per channel for display purposes only, and pseudocolours were assigned as indicated in the figure legends.

Distances, such as follicle diameters (Supplementary Figure 3C) and the thickness of the tongue epithelium and connective tissue layer (Figure 4C), were measured manually in Imaris (using the Measurement Points tool in the Slice view); the corresponding H&E measurements were made in QuPath. Lymphoid follicles, tubular structures and muscle fibres were tracked manually by following them through consecutive optical sections (Supplementary Figures 2B–D, 3D, 4C,D, 5C,D), and the Z position of each optical section along the imaging axis is indicated in the respective panels.

### Surface segmentation

The tongue epithelium (Figure 4D) was segmented in an LSFM stack of 1849 × 1849 × 5628 µm using the Imaris Surfaces module on the NR channel, with smoothing enabled (surface detail 5 µm). Epithelium was separated from the underlying tissue with the machine learning-based segmentation of the Surfaces module, which was trained by manually annotating foreground (epithelium) and background on a set of optical sections and was retrained iteratively until the prediction matched the epithelium visible in the NR signal. Splitting of touching objects was disabled, and a volume filter (≥ 1.88 × 10⁹ µm³) was applied to retain only the single continuous epithelial surface, of which the volume is reported. Muscle fibres (Figure 4G; segmented in smaller XY stacks and shown qualitatively) were segmented accordingly.

Adipocytes in the parotid gland (Figure 5G; Supplementary Figure 5E,F) were segmented in an LSFM stack of 444 × 444 × 1928 µm (0.38 mm³) using the Imaris Surfaces module on the NR channel with smoothing enabled (surface detail 3 µm) and machine learning-based segmentation, trained as described above on annotations distributed over multiple optical sections. Because adipocytes are tightly packed, touching objects were split using seed points with an estimated diameter of 25 µm. Seed points were filtered by quality (threshold set to −5.41) to obtain a single seed point per adipocyte. The resulting surfaces were then filtered by volume to remove small debris and fragments (< 4,000 µm³, corresponding to a sphere with a diameter of approximately 20 µm) and artefacts originating from tissue-free optical sections at the start of the stack (> 3.5 × 10⁵ µm³), and by sphericity (≥ 0.451).

All 1191 segmented adipocytes were used to calculate the adipocyte density (number of adipocytes per mm³ of imaged volume) and the volume fraction (summed adipocyte volume divided by the imaged volume). Size and shape measurements were restricted to the 733 adipocytes that did not touch the image border, which were identified from the Imaris statistic ‘Distance to Image Border XYZ’ after export. Border-touching adipocytes were not removed from the segmentation itself. Adipocyte diameter is reported as the equivalent spherical diameter, i.e. the diameter of a sphere with the same volume as the segmented surface and was therefore not measured directly. Sphericity was calculated by Imaris as the ratio of the surface area of a sphere with the same volume as the object to the surface area of the object, where a value of 1 indicates a perfect sphere. Nearest-neighbour distances were calculated as centroid-to-centroid distances for the 733 complete adipocytes, using all 1191 adipocytes as potential neighbours, to avoid overestimating the distance for adipocytes close to the image border.

### Statistics

Quantitative data are reported as mean ± standard deviation (SD). Calculations were performed in Microsoft Excel and R version 4.6.2 (R Foundation for Statistical Computing, Vienna, Austria).

## Acknowledgements

The authors sincerely thank the body donors and their families for their invaluable contribution to medical education and research. We thank Marjolein Verhesen (Department of Anatomy & Embryology, UM) for her help with the generation of the FFPE blocks and for providing access to the department’s equipment and expertise. We thank Gregorio Fazzi (Department of Pathology, MUMC+) for his instruction and advice on sectioning FFPE blocks and for providing access to the department’s equipment. We thank the members of the Oncometabolic Group of the Departments of Genetics & Cell Biology and Precision Medicine (Maastricht University) for useful discussions. Finally, we thank Tom Derix for his help with data visualisation.

## Author contributions

Conceptualisation: AS, FM, LDJS, ZS. Methodology: AS, FM, LDJS. Investigation: LDJS, IG, AB, EW. Resources: AS, FM, MvdH, AH. Formal analysis: LDJS, TA. Data curation: LDJS, AS, FM, ZS. Validation: MvdH, AH, FM. Visualisation: LDJS. Writing – original draft: LDJS, AS, FM. Writing – review & editing: AS, FM, LDJS, IG, EW, AH. Supervision: AS, FM. Project administration: AS, FM, LDJS. Funding acquisition: AS. All authors read and approved the final manuscript.

