## Supplementary Information for "Light-sheet fluorescence microscopy of optically cleared human head-and-neck organs: three-dimensional tissue assessment and histological correlation"

### Supplementary Figures

Supplementary Figure 1. Sampling of human head-and-neck tissues for LSFM and H&E

Supplementary Figure 2. Separate channels and 3D tracking of a lymphoid follicle and tubular structures in the palatine tonsil

Supplementary Figure 3. Separate channels, H&E overview, follicle measurements and 3D tracking of tubular structures in a cervical lymph node

Supplementary Figure 4. Separate channels, H&E overview and 3D tracking of tubular structures and muscle fibres in the tip of the tongue

Supplementary Figure 5. Separate channels, H&E overview, 3D tracking of tubular structures and adipocyte segmentation in the parotid gland

Supplementary Figure 6. Two-photon microscopy of the parotid gland compared with H&E

### Supplementary Videos

**Supplementary Video 1.** Tracking of a tubular structure through a cervical lymph node.

**Supplementary Video 2**. 3D overview of the cleared tip of the tongue..

**Supplementary Video 3**. 3D view of the ductal network within a parotid gland lobule.

### Supplementary Figures

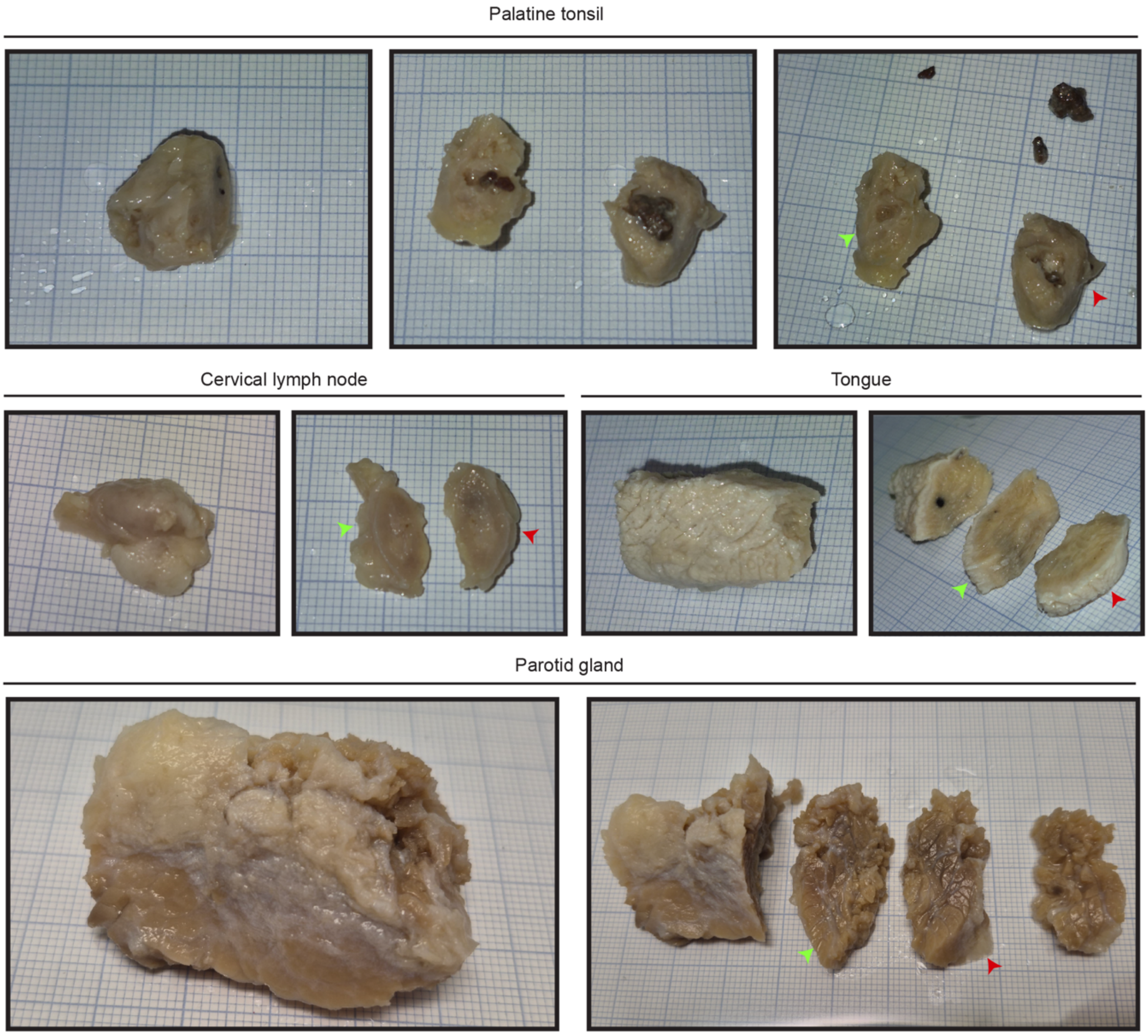


***Supplementary Figure 1. Sampling of human head-and-neck tissues for LSFM and H&E.*** *For each organ, the tissue as provided by the Department of Anatomy & Embryology is depicted on the left. The resulting tissue sections after processing of the tissue by an experienced head-and-neck surgeon are depicted on the right. The middle section of the tonsil depicts the status of the tonsil before removal of the crypt debris present as tonsillar stones. Images are captured on millimetre paper. Green arrows indicate the tissue piece selected for tissue clearing and light-sheet microscopy. Red arrows indicate the tissue piece selected for paraffin embedding and H&E. Small square: 1x1 mm.*

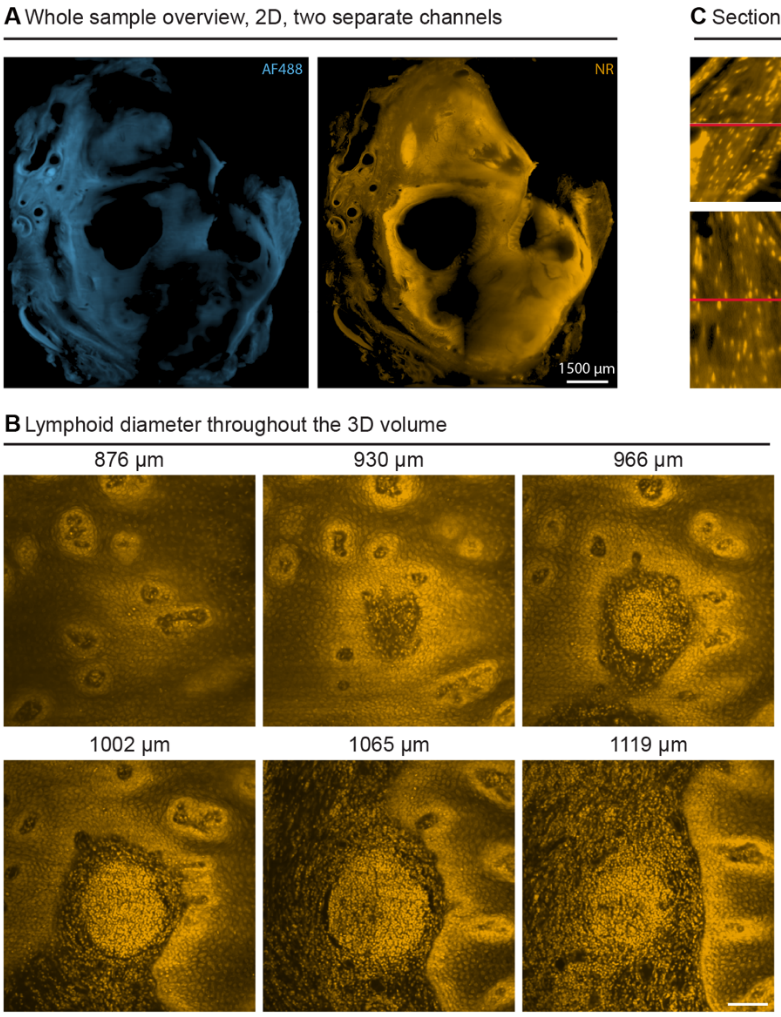


***Supplementary Figure 2. Separate channels and 3D tracking of a lymphoid follicle and tubular structures in the palatine tonsil. A)*** *Separate channels corresponding to the 2D whole sample overview image of Figure 2A.* ***B)*** *Manual tracking of a lymphoid follicle throughout different planes in the 3D volume. Z position of the optical section along the imaging axis is indicated.* ***C)*** *Orthogonal views of a tubular structure.* ***D)*** *Tracking of tubular structures through three different planes of the 3D volume. Z position of the optical section along the imaging axis is indicated. Green arrowheads show the branching of the tubular structure. Pseudocolours: autofluorescence (AF)488 (blue), NR (yellow). Scale bars and their measures are depicted in the lower right corner of each panel.*

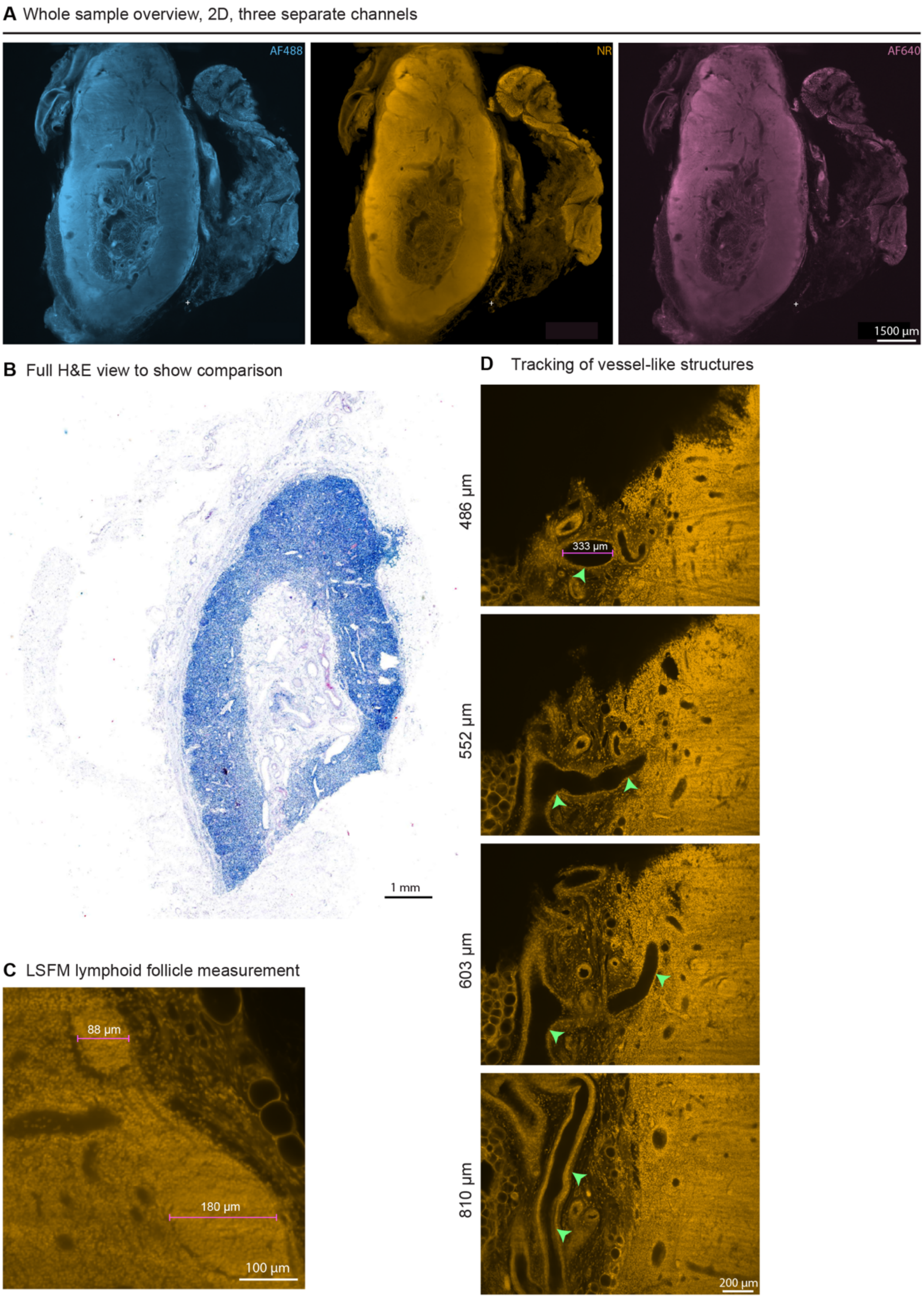


***Supplementary Figure 3. Separate channels, H&E overview, follicle measurements and 3D tracking of tubular structures in a cervical lymph node. A)*** *Separate channels corresponding to the 2D whole sample overview image of Figure 3A.* ***B)*** *Large-scale H&E overview image to illustrate macroscopic lymph node organisation.* ***C)*** *Measurement of primary lymphoid follicles imaged with LSFM.* ***D)*** *Manual tracking of tubular structures throughout different slices of the 3D LSFM volume stack to exemplify the potential pathways. Z position of the optical section along the imaging axis is indicated. Pseudocolours: autofluorescence (AF)488 (blue), NR (yellow), AF640 (magenta). Scale bars and their measures are depicted in the lower right corner of each panel.*

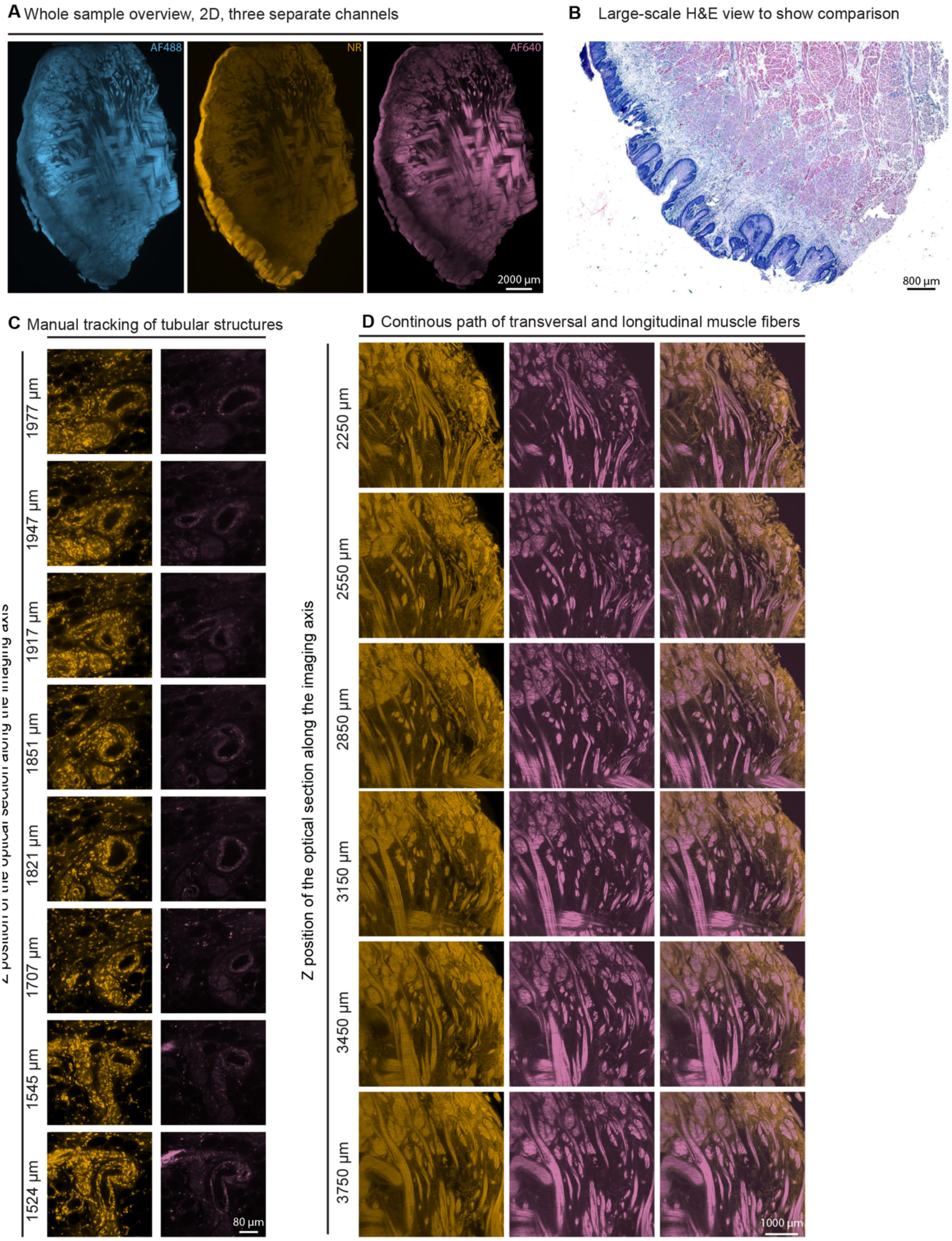


***Supplementary Figure 4. Separate channels, H&E overview and 3D tracking of tubular structures and muscle fibres in the tip of the tongue. A)*** *Separate channels corresponding to the 2D whole sample overview image of Figure 4A.* ***B)*** *Large-scale H&E overview image indicates highly diverse thickness of epithelium.* ***C, D)*** *Manual tracking of tubular structures (C) and transversal and longitudinal muscle fibres (D) throughout different slices of the 3D LSFM volume stack to exemplify the potential pathways. Z position of the optical section along the imaging axis is indicated. Pseudocolours: autofluorescence (AF)488 (blue), NR (yellow), AF640 (magenta). Scale bars and their measures are depicted in the lower right corner of each panel.*

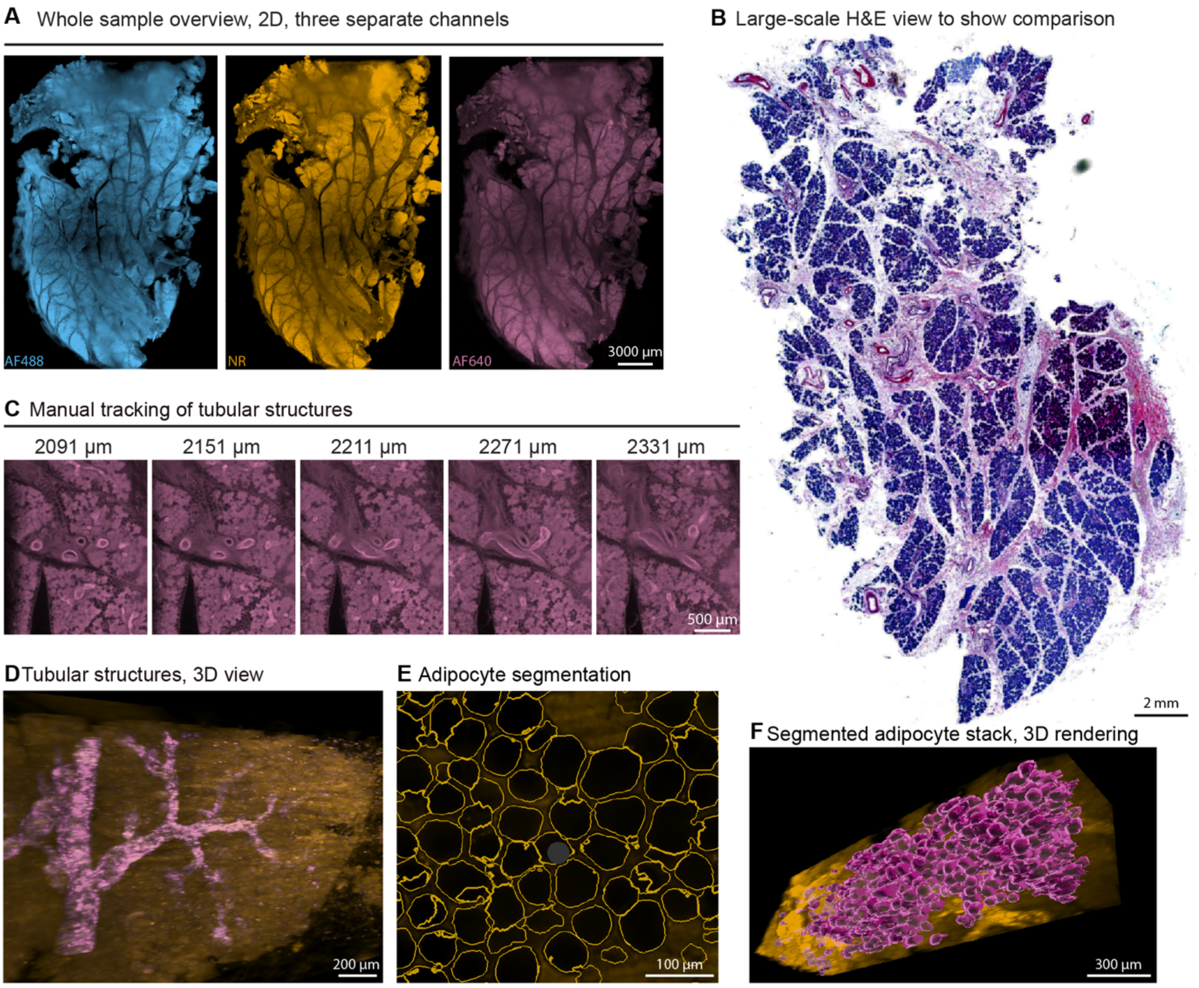


***Supplementary Figure 5. Separate channels, H&E overview, 3D tracking of tubular structures and adipocyte segmentation in the parotid gland. A)*** *Separate channels corresponding to the 2D whole sample overview image of Figure 5A.* ***B)*** *Large-scale H&E overview image illustrates macroscopic organisation of the parotid into different lobules.* ***C, D)*** *Manual tracking of tubular structures in 2D planes (C) and a 3D volume (D) to exemplify the potential pathways of the tubular structures.* ***E)*** *Example image of the adipocyte segmentation with the outline of the segmented adipocytes indicated with yellow lines.* ***F)*** *3D rendering of the complete LSFM stack (444 × 444 × 1928 µm) with the segmented adipocytes displayed as surfaces against the Neutral Red (NR) signal. Individual adipocytes are shown in purple. Pseudocolours: autofluorescence (AF)488 (blue), NR (yellow), AF640 (magenta). Scale bars and their measures are depicted in the lower right corner of each panel.*

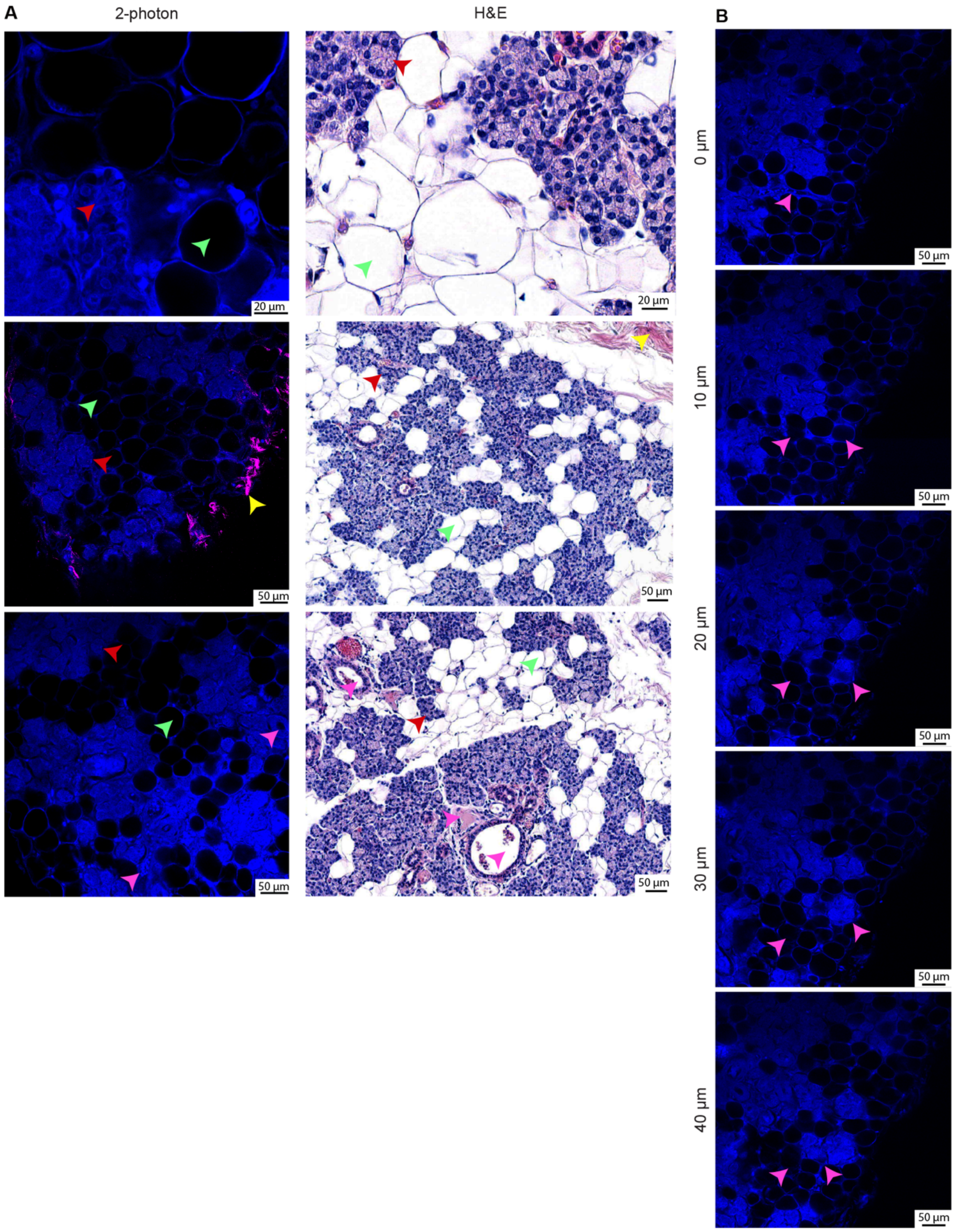


***Supplementary Figure 6. Two-photon microscopy of the parotid gland compared with H&E. A)*** *Comparison of two-photon imaging with H&E to show comparable structures like adipocytes (green arrowheads), acini (red arrowheads), connective tissue (yellow arrowheads) and tubular structures (pink arrowheads).* ***B)*** *Two-photon image stack. Pink arrowheads show changing structures. Z position of the optical section along the imaging axis is indicated. Pseudocolours: DAPI (blue), second harmonic generation (magenta). Scale bars and their measures are depicted in the lower right corner of each panel.*

### Supplementary Videos

**Supplementary Video 1.** Tracking of a tubular structure through a cervical lymph node. A tubular structure, possibly a lymph or blood vessel, is followed through consecutive optical sections of the LSFM volume over approximately 1500 µm. Pseudocolour: neutral red (yellow). Scale bar: 300 µm.

**Supplementary Video 2**. 3D overview of the cleared tip of the tongue. Volume rendering of the complete LSFM dataset of the tongue tip (22 × 13 × 7.8 mm; 2.2 $cm^3$), showing the epithelium, connective tissue layer and the network of intrinsic muscle fibres. Pseudocolours: autofluorescence (AF)488 (blue), neutral red (yellow), AF640 (magenta). Scale bar: 3000 µm.

**Supplementary Video 3**. 3D view of the ductal network within a parotid gland lobule. Volume rendering of a single lobule, showing the organised intralobular network of tubular structures. Pseudocolours: AF640 (magenta). Scale bar: 700 µm.
